# Interoperable and scalable data analysis with microservices: Applications in Metabolomics

**DOI:** 10.1101/213603

**Authors:** Payam Emami Khoonsari, Pablo Moreno, Sven Bergmann, Joachim Burman, Marco Capuccini, Matteo Carone, Marta Cascante, Pedro de Atauri, Carles Foguet, Alejandra Gonzalez-Beltran, Thomas Hankemeier, Kenneth Haug, Sijin He, Stephanie Herman, David Johnson, Namrata Kale, Anders Larsson, Steffen Neumann, Kristian Peters, Luca Pireddu, Philippe Rocca-Serra, Pierrick Roger, Rico Rueedi, Christoph Ruttkies, Noureddin Sadawi, Reza M Salek, Susanna-Assunta Sansone, Daniel Schober, Vitaly Selivanov, Etienne A. Thevenot, Michael van Vliet, Gianluigi Zanetti, Christoph Steinbeck, Kim Kultima, Ola Spjuth

**Affiliations:** Department of Medical Sciences, Clinical Chemistry, Uppsala University, Uppsala, Sweden; European Molecular Biology Laboratory, European Bioinformatics Institute (EMBL-EBI), Cambridge, United Kingdom; Department of Computational Biology, University of Lausanne, Lausanne, Switzerland; Swiss Institute of Bioinformatics, Lausanne, Switzerland; Department of Neuroscience, Uppsala University, Uppsala, Sweden; Department of Information Technology, Uppsala, Sweden; Department of Pharmaceutical Biosciences, Uppsala University, Uppsala, Sweden; Department of Biochemistry and Molecular Biomedicine, Faculty of Biology, Universitat de Barcelona, Barcelona, Spain; Institute of Biomedicine of the Universitat de Barcelona (IBUB) and Associated Unit to CSIC, Barcelona, Spain; Oxford e-Research Centre, Department of Engineering Science, University of Oxford, Oxford, United Kingdom; Division of Analytical Biosciences, Leiden Academic Centre for Drug Research, Leiden University, Leiden, The Netherlands; National Bioinformatics Infrastructure Sweden, Uppsala University, Uppsala, Sweden; Department of Stress and Developmental Biology, Leibniz Institute of Plant Biochemistry, Halle, Germany; German Centre for Integrative Biodiversity Research (iDiv), Halle-Jena-Leipzig, Germany; CRS4: Center for Advanced Studies, Research and Development in Sardinia, Pula, Italy; CEA, LIST, Laboratory for Data Analysis and Systems' Intelligence, MetaboHUB, Gif-sur-Yvette, France; Faculty of Medicine, Department of Surgery & Cancer, Imperial College London, London, United Kingdom; International Agency for Research on Cancer 150 cours Albert Thomas 69372 Lyon CEDEX 08, France; Friedrich-Schiller-University, Jena, Germany

## Abstract

Developing a robust and performant data analysis workflow that integrates all necessary components whilst still being able to scale over multiple compute nodes is a challenging task. We introduce a generic method based on the microservice architecture, where software tools are encapsulated as Docker containers that can be connected into scientific workflows and executed in parallel using the Kubernetes container orchestrator. The access point is a virtual research environment which can be launched on-demand on cloud resources and desktop computers. IT-expertise requirements on the user side are kept to a minimum, and established workflows can be re-used effortlessly by any novice user. We validate our method in the field of metabolomics on two mass spectrometry studies, one nuclear magnetic resonance spectroscopy study and one fluxomics study, showing that the method scales dynamically with increasing availability of computational resources. We achieved a complete integration of the major software suites resulting in the first turn-key workflow encompassing all steps for mass-spectrometry-based metabolomics including preprocessing, multivariate statistics, and metabolite identification. Microservices is a generic methodology that can serve any scientific discipline and opens up for new types of large-scale integrative science.

## INTRODUCTION

Biology is becoming data-intensive as high throughput experiments in genomics or metabolomics are rapidly generating data sets of massive volume and complexity (1, 2), posing a fundamental challenge on large scale data analytics.

Currently, the most common large-scale computational infrastructures in science are shared High-Performance Computing (HPC) systems. Such systems are usually designed primarily to support computationally intensive batch jobs – e.g., for the simulation of physical processes – and are managed by specialized system administrators. This model leads to rigid constraints on the way these resources can be used. For instance, the installation of software must undergo approval and may be restricted, which contrasts with the needs in the analysis where a multitude of software components of various versions – and their dependencies – are needed, and where these need to be continuously updated.

Cloud computing offers a compelling alternative to shared HPC systems, with the possibility to instantiate and configure on-demand resources such as virtual computers, networks, and storage, together with operating systems and software tools. Users only pay for the time the virtual resources are used, and when they are no longer needed they can be released and incur no further costs for usage or ownership.

Along with infrastructure provisioning, software provisioning – i.e., installing and configuring software for users – has also advanced. Consider, for instance, containerization (3), which allows entire applications with their dependencies to be packaged, shipped and run on a computer but isolated from one another in a way analogous to virtual machines, yet much more efficiently. Containers are more compact, and since they share the same operating system kernel, they are fast to start and stop and incur little overhead in execution. These traits make them an ideal solution to implement lightweight *microservices*, a software engineering methodology in which complex applications are divided into a collection of smaller, loosely coupled components that communicate over a network (4). Microservices share many properties with traditional always-on web services found on the Internet, but microservices are generally smaller, portable and can be started on-demand within a separate computing environment. Another important feature of microservices is that they have a technology-agnostic communication protocol, and hence can serve as building blocks that can be combined and reused in multiple ways (5).

Microservices are highly suitable to run in elastic cloud environments that can dynamically grow or shrink on demand, enabling applications to be scaled-up by simply starting multiple parallel instances of the same service. However, to achieve effective scalability a system needs to be appropriately sectioned into microservice components and the data to be exchanged between the microservices needs to be defined for maximum efficiency–both being challenging tasks.

One of the omics fields that faces challenges by data growth is metabolomics which measures the occurrence, concentrations and changes of small molecules (metabolites) in organisms, organs, tissues, cells and cellular compartments. Metabolite abundances are assayed in the context of environmental or dietary changes, disease or other conditions (6). Metabolomics is, as most other omics technologies, characterized by the use of high-throughput experiments performed using a variety of spectroscopic methods such as Mass Spectrometry (MS) and Nuclear Magnetic Resonance (NMR) that produce large amounts of data (7). With increasing data size and number of samples, the analysis process becomes intractable for desktop computers due to requirements on compute cores, memory, storage etc. As a result, large-scale computing infrastructures have become important components in scientific projects (8). Moreover, making use of such complex computing resources in an analysis workflow presents its own challenges, including achieving efficient job parallelism and scheduling as well as error handling (9). In addition, configuring the necessary software tools and chaining them together into a complete re-runnable analysis workflow commonly requires substantial IT-expertise, while creating portable and fault-tolerant workflows with a robust audit trail is even more difficult. Metabolomics has already benefited from cloud-based systems such as XCMS ONLINE (10), Chorus (chorusproject.org) and The Metabolomics Workbench (www.metabolomicsworkbench.org) which provide virtual environments that scale with computational demands. However, these applications provide limited flexibility in terms of incorporating and maintaining tools as well as constructing and using customizable workflows.

In this manuscript, we present a method which uses components for metabolomics data analysis encapsulated as microservices and connected into computational workflows to provide complete, ready-to-run, reproducible data analysis solutions that can be easily deployed on desktop computers as well as public and private clouds. Our approach requires virtually no involvement in the setup of computational infrastructure and no special IT skills from the user. We validate the method on four metabolomics studies and show that it enables scalable and interoperable data analysis.

## Material and methods

### Microservices

A detailed description of the methods is present in supplementary method S1. Briefly, in order to construct a microservice architecture for metabolomics we used Docker (11) (https://www.docker.com/) containers to encapsulate software tools. Tools are developed as open source and are available in a public repository such as GitHub (https://github.com/). The containers are built and tested on a Jenkins continuous integration (CI) server (https://jenkins.io/).

### Virtual Research Environment (VRE)

We developed a Virtual Research Environment (VRE) which uses Kubernetes (https://kubernetes.io/) for orchestration of the containers, including initialisation and scaling of jobs based on containers, abstractions to file system access for running containers, exposure of services, as well as rescheduling of failed jobs and long running services. In addition, to enable convenient instantiation of a complete virtual infrastructure, we developed KubeNow (https://github.com/kubenow/KubeNow) which includes instantiation of compute nodes, shared file system storage, networks, configure DNS, operating system, container implementation and orchestration tools on a local computer or server. In order to deploy applications, we used two main classes of services: long-lasting services, and compute jobs. Long-lasting services were used for applications such as the user interface whereas compute jobs were used to perform temporary functions in data processing.

### Demonstrators

We validated our method in the field of metabolomics using four demonstrators. Demonstrators 1 and 2 showcase scalability and interoperability of our microservice-based architecture whereas Demonstrators 3 and 4 exemplify flexibility to account for new application domains, showing the architecture is domain-agnostic.

#### Demonstrator 1: Scalability of microservices in a cloud environment

The objective of this analysis was to demonstrate the computational scalability of an existing workflow on a large dataset (MetaboLights (12) ID: MTBLS233, http://www.ebi.ac.uk/metabolights/MTBLS233 (13)). The experiment includes 528 mass spectrometry samples from whole cell lysates of human renal proximal tubule cells that were pre-processed through a five-step workflow (consisting of peak picking, feature finding, linking, file filtering and exporting) using the OpenMS software (14). This preprocessing workflow was reimplemented using Docker containers and run using the Luigi workflow engine. Scalability of concurrent running tools (on 40 Luigi workers, each worker receives tasks from the scheduler and executes them) was measured using weak scaling efficiency (WSE), where the workload assigned to each worker stays constant and additional workers are used to solve a larger total problem.

#### Demonstrator 2: Interoperability of microservices

The objective of this analysis was to demonstrate interoperability as well as to present a real-world scenario in which patients’ data are processed using a microservices-based platform. We used a dataset consisting of 37 clinical cerebrospinal fluid (CSF) samples including thirteen relapsing-remitting multiple sclerosis (RRMS) patients and 14 secondary progressive multiple sclerosis (SPMS) patients as well as 10 non-multiple sclerosis controls. 26 quality controls (19 blank and 7 dilution series samples) were also added to the experiment. In addition, 8 pooled CSF samples containing MS/MS data were included in the experiment for improving identification (MetaboLights ID: MTBLS558, http://www.ebi.ac.uk/metabolights/MTBLS558). The samples were processed and analysed on the Galaxy platform (15), running in a the VRE behind the Uppsala University Hospital firewall to be compliant with local ELSI (Ethics, Legal, Social implications) regulations.

#### Demonstrator 3:1D NMR-analysis workflow

The purpose of this demonstrator was to highlight the fact that the microservice architecture is indeed domain-agnostic and is not limited to a particular assay technology. This NMR-based metabolomics study was originally performed by Salek et al.(16) on urine of type 2 diabetes mellitus (T2DM) patients and controls (MetaboLights ID: MTBLS1, http://www.ebi.ac.uk/metabolights/MTBLS1). In total, 132 samples (48 T2DM and 84 controls) were processed using a Galaxy workflow performing conversion, preprocessing, multivariate data analysis and result visualization.

#### Demonstrator 4: Start-to-end fluxomics workflow

The purpose of this demonstrator was to show the integrated use of separately developed tools covering subsequent steps of the study of metabolic fluxes based on ^13^C stable isotope-resolved metabolomics (SIRM)(17-19). Here we implemented the analysis of flux distributions in HUVEC cells under hypoxia (MetaboLights ID: MTBLS412, http://www.ebi.ac.uk/metabolights/MTBLS412), from raw mass spectra contained in netCDF files, using a workflow implemented in Galaxy including reading and extraction of the data, correcting the evaluated mass spectra for natural isotopes and computing steady-state distribution of ^13^C label as function of steady-state flux distribution.

### Availability

The PhenoMeNal consortium maintains a web portal (https://portal.phenomenal-h2020.eu) providing a GUI for launching VREs using KubeNow (20) on a selection of the largest public cloud providers, including Amazon Web Services, Microsoft Azure and Google Cloud Platform, or on private OpenStack-based installations. The containers provisioned by PhenoMeNal comprise tools built as open source software that are available in a public repository such as GitHub, and are subject to continuous integration testing. The containers that satisfy testing criteria are pushed to a public container repository, and containers that are included in stable VRE releases are also pushed to Biocontainers (21).

## Results

We developed a VRE based on a microservices architecture encapsulating a large suite of software tools for performing metabolomics data analysis (See Table S1). Scientists can interact with the microservices programmatically via an Application Programming Interface (API) or via a web-based graphical user interface (GUI), as illustrated in Figure 1. To connect microservices into computational workflows, the two frameworks Galaxy (15) and Luigi (https://github.com/spotify/luigi) were adapted to execute jobs on Kubernetes. Galaxy is a web-based interface for individual tools and allows users to share workflows, analysis histories and result data sets. Luigi on the other hand focuses on scheduled execution, monitoring, visualization and the implicit dependency resolution of tasks (22). These basic infrastructure services, together with the Jupyter notebook (23) interactive programming environment, are deployed as long-running services in the VRE, whereas the other analysis tools are deployed as transient compute jobs to be used on-demand. System and client applications were developed for launching the VRE on desktop computers, public and private cloud providers, automating all steps required to instantiate the virtual infrastructures.

**Figure 1:**
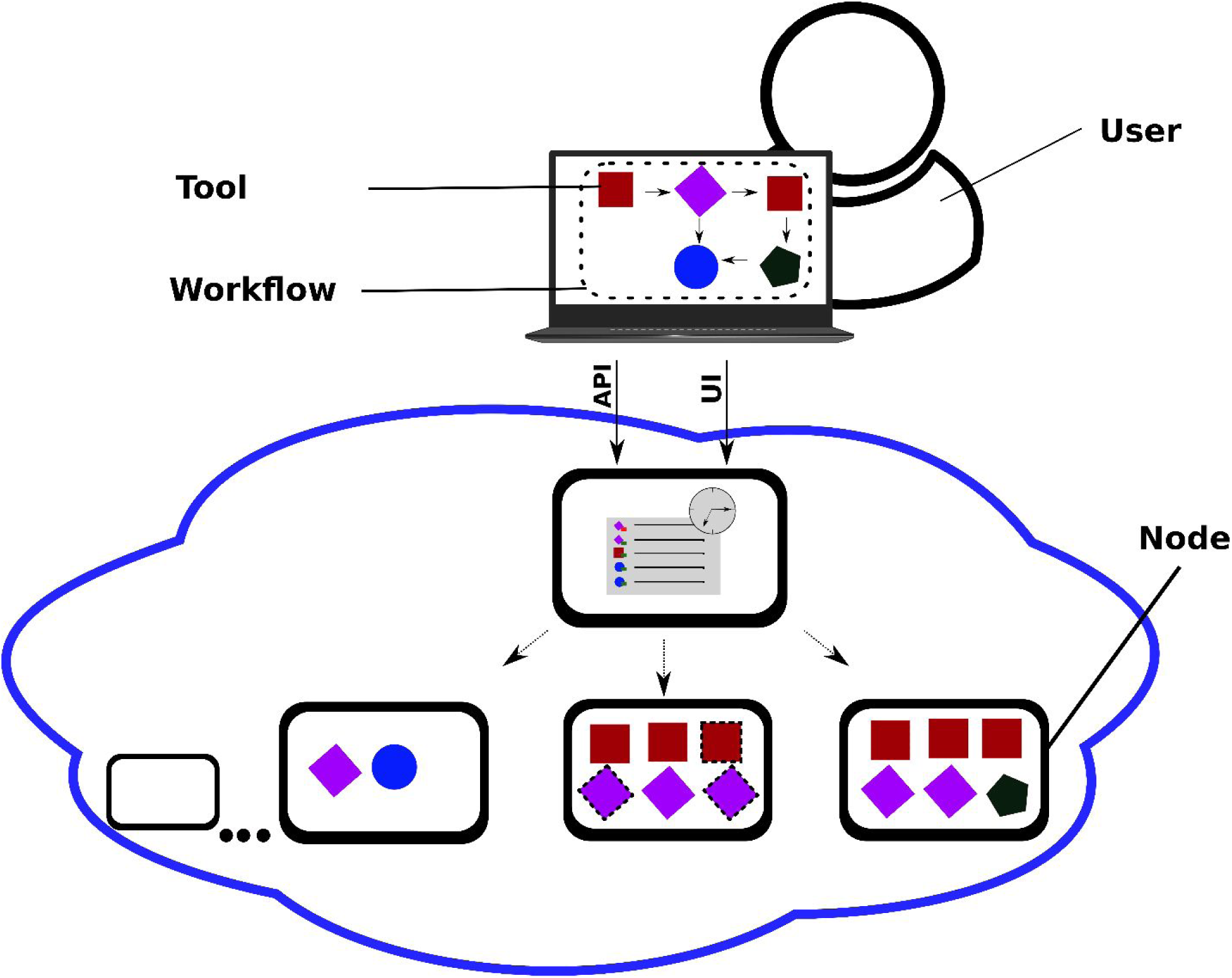
Overview of the components in a microservices-based framework. Complex applications are divided into smaller, focused and well-defined (micro-) services. These services are independently deployable and can communicate with each other, which allows to couple them into complex task pipelines, i.e. data processing workflows. The user can interact with the framework programmatically via an Application Program Interface (API) or via a graphical user interface (GUI) to construct or run workflows of different services, which are executed independently. Multiple instances of services can be launched to execute tasks in parallel, which effectively can be used to scale analysis over multiple compute nodes. When run in an elastic cloud environment, virtual resources can be added or removed depending on the computational requirements.

### Demonstrator 1: Scalability of microservices in a cloud environment

The Diagram of scalability-testing on the metabolomics dataset is illustrated in Figure 2. The analysis resulted to WSE of 88% with an execution time of approximately four hours (online methods, Figure S2), compared with the ideal case of 100% where linear scaling is achieved if the run time stays constant while the workload is increased. In addition, the final result of the workflow (online methods, Figure S3) was identical to that presented by the original MTBLS233 study (Ranninger et al.(13)) in negative ionization mode. However, in the positive ionization mode, one m/z feature was found in a different group (m/z range 400-1000) than it was originally reported by Ranninger et al. (m/z range 200-400).

**Figure 2.**
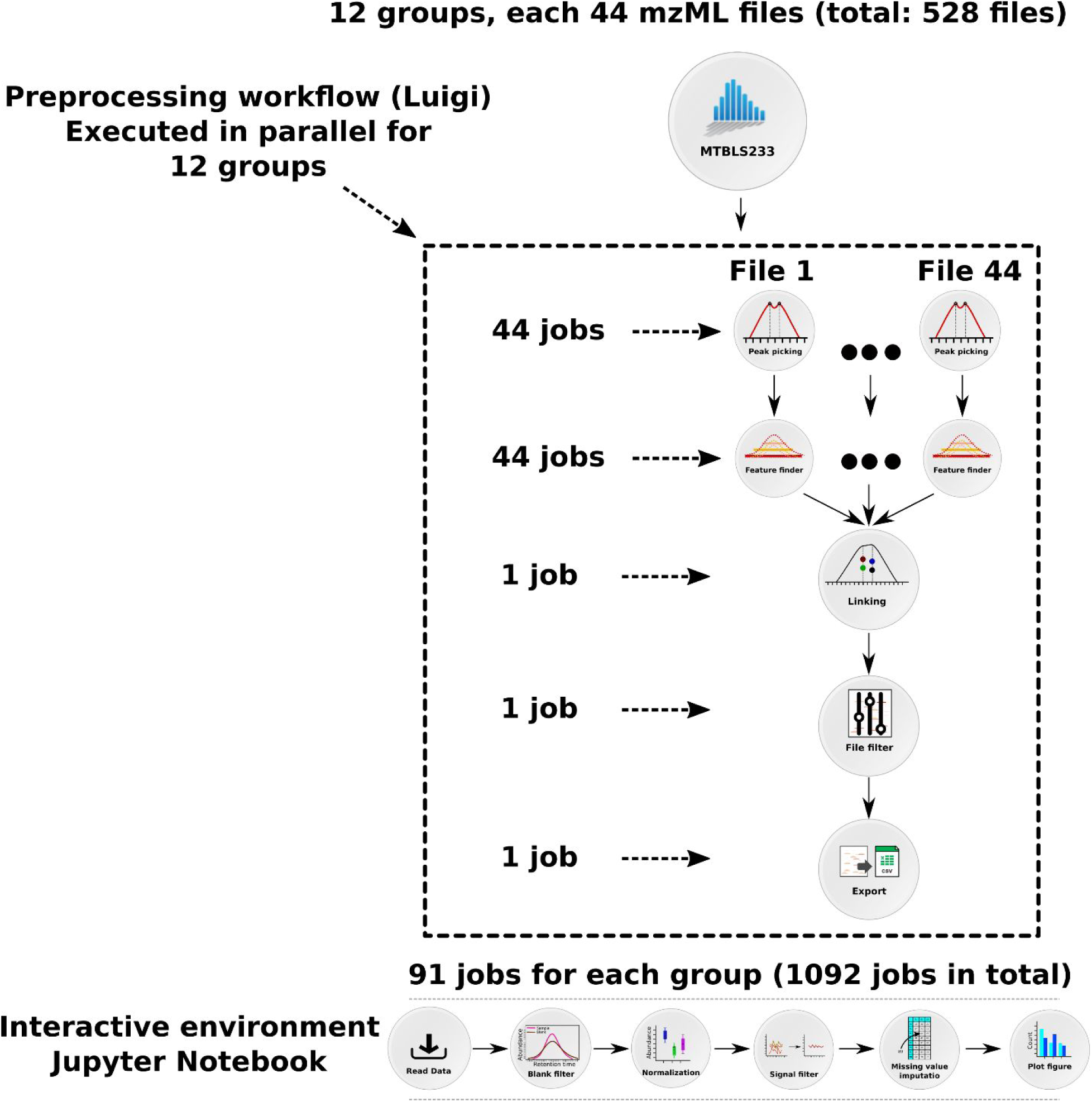
Diagram of scalability-testing on a metabolomics dataset (MetaboLights ID: MTBLS233) in Demonstrator 1 to illustrate the scalability of a microservice approach. The preprocessing workflow is composed of 5 OpenMS tasks that were run in parallel over the 12 groups in the dataset using the Luigi workflow system. The first two tasks, peak picking (528 tasks) and feature finding (528 tasks), are trivially parallelizable, hence they were run concurrently for each sample. The subsequent feature linking task needs to process all of the samples in a group at the same time, therefore 12 of these tasks were run in parallel. In order to maximize the parallelism, each feature linker container (microservice) was run on 2 CPUs. Feature linking produces a single file for each group, that can be processed independently by the last two tasks: file filter (12 tasks) and text exporter (12 tasks), resulting in total of 1092 tasks. The downstream analysis consisted of 6 tasks that were carried out in a Jupyter Notebook. Briefly, the output of preprocessing steps was imported into R and the unstable signals were filtered out. The missing values were imputed and the resulting number of features were plotted.

### Demonstrator 2: Interoperability of microservices

We developed a start to end workflow for pre-processing and statistical analysis of LC-MS metabolomics data (Figure 3). The workflow allows seamless integration of six major metabolomics data analysis components (26 steps) each was already implemented in independent software suites: noise reduction and filtering (OpenMS (24)), quantification, alignment and matching (XCMS (25)), filtering of biological non-relevant signals (R), annotation of signals (CAMERA (26)), identification (MetFrag (27)), statistics (Workflow4Metabolomics (28)). The result of the workflow (multivariate analysis) showed a clear difference in the metabolic constitution between the three disease groups of RRMS, SPMS and non-multiple sclerosis controls (Figure 4A). In addition, the univariate analysis resulted in a total of three metabolites being significantly altered (p<0.05) between multiple sclerosis subtypes and control samples, namely alanyltryptophan and indoleacetic acid with higher and linoleoyl ethanolamide with lower abundance in both RRMS and SPMS compared to controls (Figure 4B).

**Figure 3.**
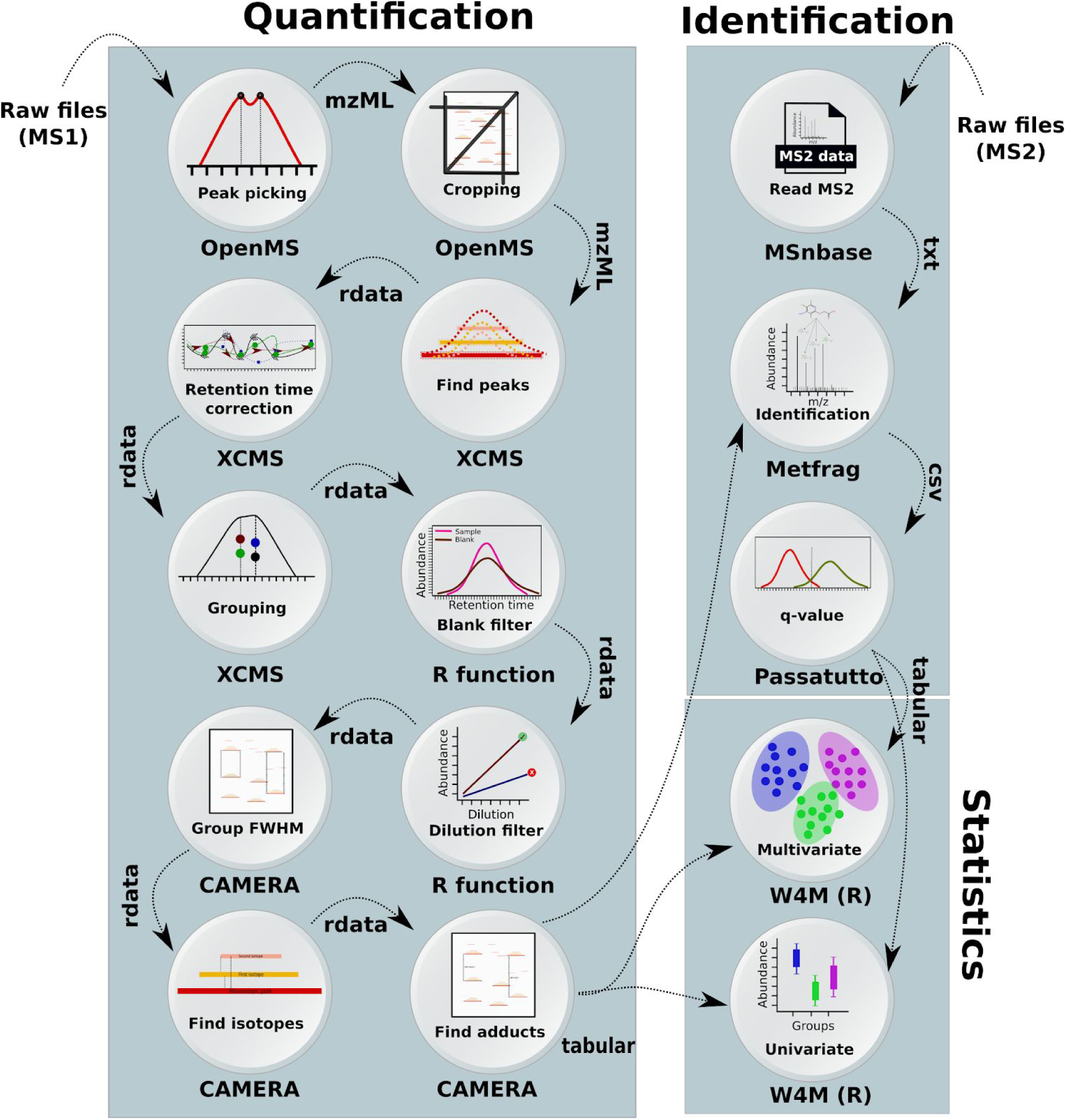
Overview of the workflow used to process multiple-sclerosis samples in Demonstrator 2, where a workflow was composed of the microservices using the Galaxy system. The data was centroided and limited to a specific mass over charge (m/z) range using OpenMS tools. The mass traces quantification and retention time correction was done via XCMS(25). Unstable signals were filtered out based on the blank and dilution series samples using an in-house function (implemented in R). Annotation of the peaks was performed using CAMERA(26). To perform the metabolite identification, the tandem spectra from the MS/MS samples in mzML format were extracted using MSnbase and passed to MetFrag. The MetFrag scores were converted to q-values using Passatutto software. The result of identification and quantification were used in “Multivariate” and “Univariate” containers from Workflow4Metabolomics(28) to perform Partial Least Squares Discriminant Analysis (PLS-DA)(49).

**Figure 4.**
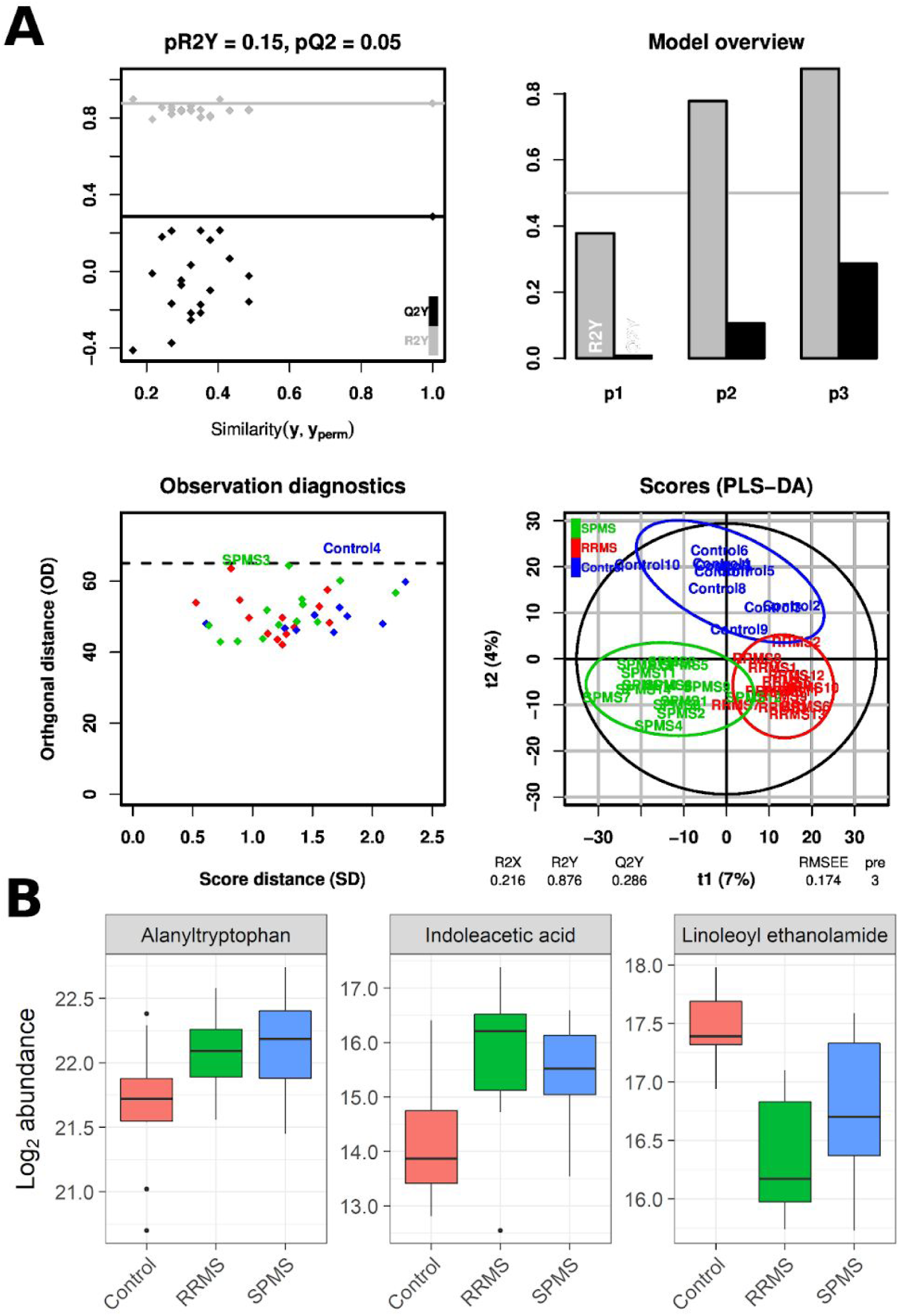
The results from analysis of multiple sclerosis data in Demonstrator 2, presenting new scientifically useful biomedical knowledge. A) The PLS-DA results suggest that the metabolite distribution in the RRMS and SPMS samples are different to controls. B) Three metabolites were identified as differentially regulated between multiple sclerosis subtypes and control samples, namely Alanyltryptophan and Indoleacetic acid with higher and Linoleoyl ethanolamide with lower abundance in both RRMS and SPMS compared to controls. Abbr., RRMS: relapsing-remitting multiple sclerosis, SPMS: secondary progressive multiple sclerosis.

### Demonstrators 3 and 4: Domain agnosticity (NMR and fluxomics workflows)

We developed a workflow for analysis of 1D NMR data. The workflow consisted of automatic downloading NMR vendor data (and metadata) from MetaboLights database followed by format standardisation, spectral processing and statistical analysis. We processed a NMR dataset (demonstrator 3) resulting to quantification of a total of 726 features which were used to perform Orthogonal Projections to Latent Structures Discriminant Analysis (OPLS-DA). This resulted in a clear separation between T2DM and controls (Figure 5), similar to that of previous findings (29). Lastly, we designed a workflow for analyzing metabolite metabolic fluxes. The workflow integrated four main steps including data extraction, data correction, calculation of flux distribution and visualisation. Using this workflow (Figure 6), we achieved detailed description of the magnitudes of the fluxes through the reactions accounting for glycolysis and pentose phosphate pathway.

**Figure 5.**
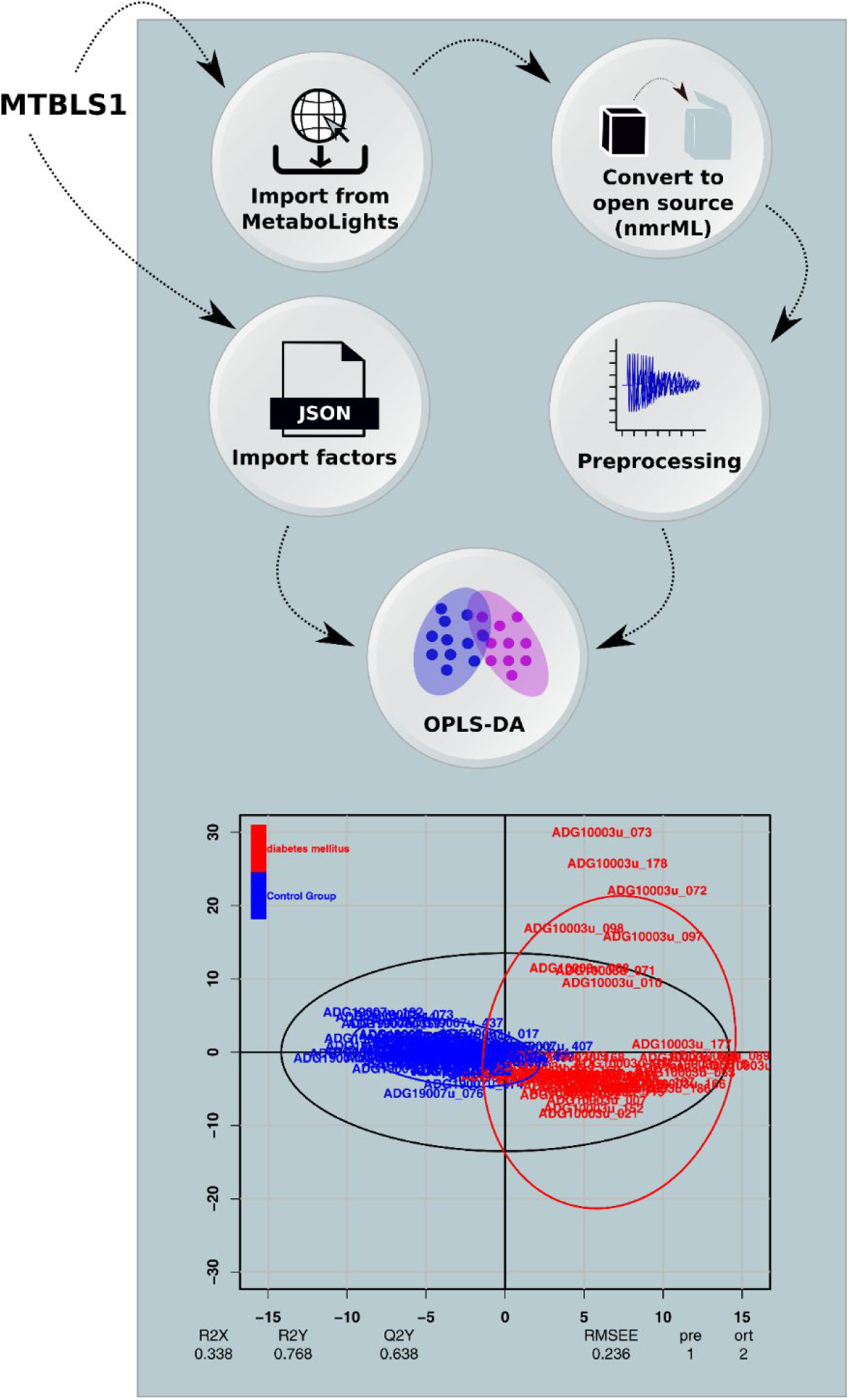
Overview of the NMR workflow in Demonstrator 3. The raw NMR data and experimental metadata (ISATab) was automatically imported from the Metabolights database and converted to open source nmrML format. The preprocessing was performed using the rnmr1d package part of nmrprocflow (50) tools. All study factors were imported from MetaboLights and were fed to the multivariate node to perform an OPLS-DA.

**Figure 6:**
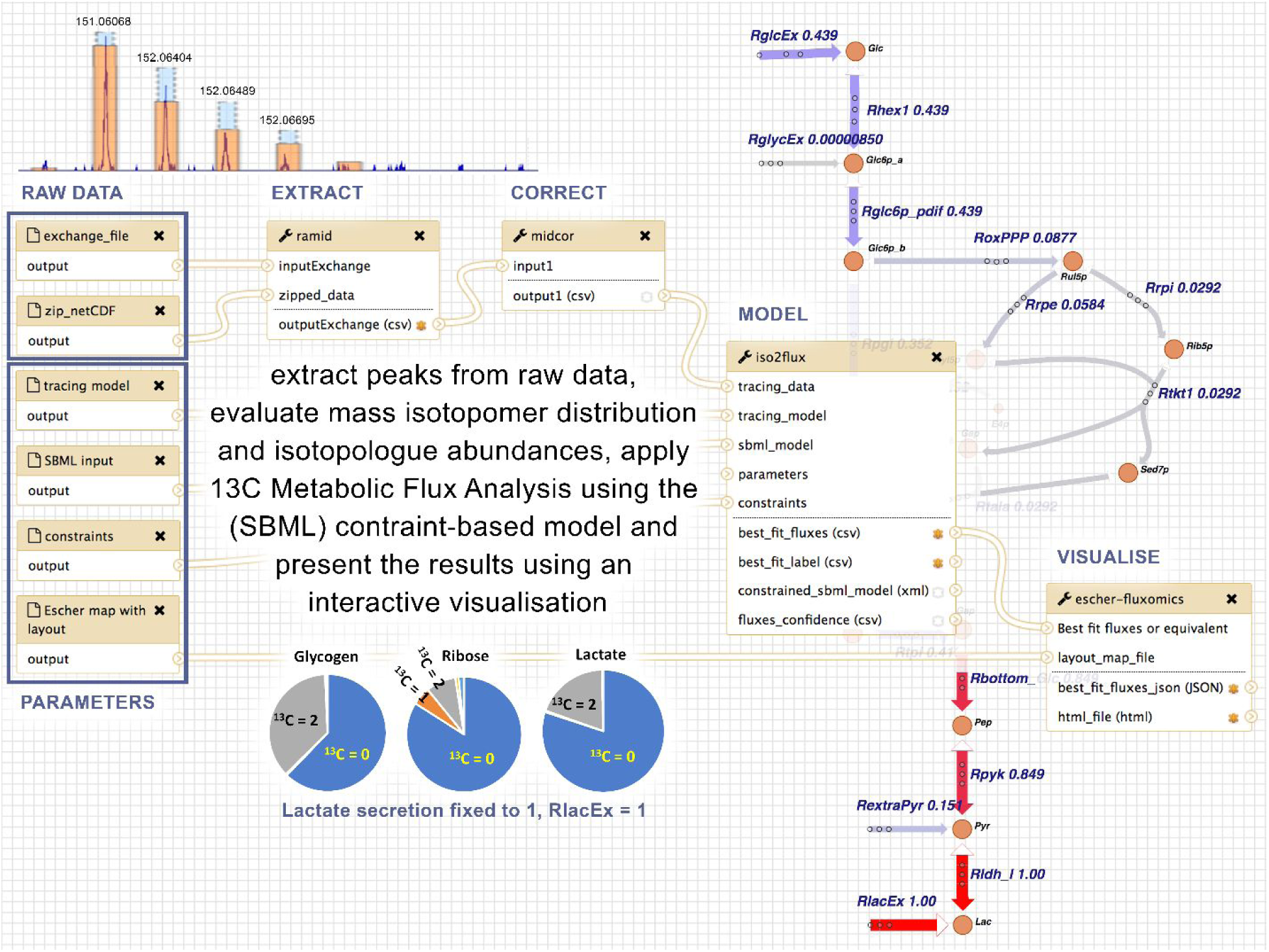
Overview of the workflow for fluxomics, with Ramid, Midcor, Iso2Flux and Escher-fluxomics tools supporting subsequent steps of the analysis. The example refers to HUVEC cells incubated in the presence of [1,2-^13^C_2_]glucose and label (^13^C) propagation to glycogen, RNA ribose and lactate measured by mass spectrometry. Ramid reads the raw netCDF files, corrects baseline and extracts the peak intensities. The resulting peak intensities are corrected (natural abundance, overlapping peaks) by Midcor, which provides isotopologue abundances. Isotopologue abundances, together with a model description (SBML model, tracing data, constraints), are used by Iso2Flux to provide flux distributions through glycolysis and pentose-phosphate pathways, which are shown as numerical values associated to a metabolic scheme of the model by the Escher-fluxomics tool.

## Discussion

Implementing the different tools and processing steps of a data analysis workflow as separate services that are made available over a network was in the spotlight in the early 2000’s (30) as service-oriented architectures (SOA) in science. At that time, web services were commonly deployed on physical hardware and exposed and consumed publicly over the internet. However, it soon became evident that this architecture did not fulfill its promises as it did not scale well from a computational perspective. In addition, the web services were not portable and mirroring them was complicated (if at all possible). Furthermore, API changes and frequent services outage made it frustrating to connect them into functioning computational workflows. Ultimately, the ability to replicate an analysis on local and remote hardware (such as a computer cluster) was very difficult due to heterogeneity in the computing environments.

At first sight microservices might seem similar to abovementioned SOA web services, but microservices can with great benefit be executed in virtual environments (abstracting over OS and hardware architectures) in such a way that they are only instantiated and executed on-demand, and then terminated when they are no longer needed. This makes such virtual environments inherently portable and they can be launched on demand on different platforms (e.g., a laptop, a powerful physical server or an elastic cloud environment). A key aspect is that workflows of microservices are still executed identically, agnostic of the underlying hardware platform. Container-based microservices provide a wide flexibility in terms of versioning, allowing the execution of newer and older versions of each container as needed for reproducibility. Since all software dependencies are encompassed within the container, which is versioned, the risk of workflow failure due to API changes is minimized. An orchestration framework such as Kubernetes further allows for managing errors in execution and transparently handles the restarting of services. Hence, technology has caught up with service-oriented science, and microservices have taken the methodology to the next level, alleviating many of the previous problems related to scalability, portability and interoperability of software tools. This is advantageous in the context of omics analysis, which produces multidimensional data sets reaching beyond gigabytes, on into terabytes, leading to ever-increasing demand on processing performance (1, 2).

In Demonstrator 1, we showed that microservices enable highly efficient and scalable data analyses by executing individual modules in parallel, and that they effectively harmonize with on-demand elasticity of the cloud computing paradigm. The reached scaling efficiency of ~88% indicates remarkable performance achieved on generic cloud providers. Furthermore, although our results in positive ionization model was slightly different to that of Ranninger et al. (13), the results of our analysis were reproducible regardless of the platform used to perform the computations, indicating a level of replicability of study results and reusability of workflows in the analysis that - to the best of our knowledge - has never been reported before in metabolomics data analysis.

In addition to the fundamental demand for high performance, the increased throughput and complexity of omics experiments has led to a large number of sophisticated computational tools (31), which in turn necessitates integrative workflow engines (32). In order to integrate new tools in such workflow engines, compatibility of the target environment, tools and APIs needs to be considered (32). Containerization facilitates this by providing a platform-independent virtual environment for developing and running the individual tools. However, the problem of compatibility between tools/APIs and data formats remains and needs to be tackled by international consortia (e.g., strictly adhering to FAIR Data Principles (33)). We also overcome the currently non-trivial task of instantiating the complete microservice environment through a web portal that allows for convenient deployment of the VRE on public cloud providers. Moreover, using this web portal, microservices and VREs can be deployed on a trusted private cloud instance or a local physical server on an internal network, such as within a hospital network, allowing for levels of isolation and avoiding transfer of data across untrusted networks which often are requirements in the analysis of sensitive data. This was exemplified in Demonstrator 2, where a complete start-to-end workflow was run on the Galaxy platform on a secure server at Uppsala University Hospital, Sweden, leading to the identification of novel disease fingerprints in the CSF metabolome of RRMS and SPMS patients. It is worth mentioning that the selected metabolites were part of the tryptophan metabolism (alanyltryptophan and indoleacetic acid) and endocannabinoids (linoleoyl ethanolamide), both of which have been previously implicated in multiple sclerosis (34-39). However, since the cross-validated predictive performance (Q2Y = 0.286) is not much higher than some of the models generated after random permutation of the response (Figure 4A), the quality of the model needs to be confirmed in a future study on an independent cohort of larger size.

The microservice architecture is domain-agnostic and not limited to a particular assay technology, i.e. mass spectrometry. This was showcased in Demonstrator 3 and 4, where an automated 1D NMR workflow and calculation of flux distributions (derived from the application of stable isotope resolved metabolomics) were performed. In Demonstrator 3, we showed that the pattern of the metabolite expression is different between type 2 diabetic and healthy controls, and that a large number of metabolites contribute to such separation. In Demonstrator 4, we showed a high rate of glycolysis in cells cultured in hypoxia, which is consistent with the one expected for endothelial cells (40) and with how these cells maintain energy in low oxygen environments and without oxidative phosphorylation (41, 42). These two examples further show that complex workflows can be applied with minimal effort on other studies (i.e. simply by providing a MetaboLights accession number), leading to the capability to re-analyze data and compare the results with the original publication findings. Furthermore, it demonstrates the value of standardised dataset descriptions such as nmrML (43) and ISA format (44, 45) for representing NMR based studies, as well as the potential of the VRE to foster reproducibility.

While microservices are not confined to metabolomics and generally applicable to a large variety of applications, there are some important implications and limitations of the method. Firstly, tools need to be containerized in order to operate in the environment. This is however not particularly complex, and an increasing number of developers provide containerized versions of their tools on public container repositories such as Dockerhub or Biocontainers (5). Secondly, uploading data to a cloud-based system can take a considerable amount of time, and having to re-do this every time a VRE is instantiated can be time-consuming. This can be alleviated by using persistent storage on a cloud resource, but the availability of such storage varies between different cloud providers. Further, the storage system can become a bottleneck when many services try to access a shared storage. We observe that using a distributed storage system with multiple storage nodes can drastically increase performance, and the PhenoMeNal VRE comes with a distributed storage system by default. When using a workflow system to orchestrate the microservices, stability and scalability are inherently dependent on the workflow system’s job runner. Workflow execution is dependent on the underlying workflow engine, and we observed that a large number of outputs can make the Galaxy engine unresponsive, whereas the Luigi engine did not have these shortcomings. With clouds and microservices maturing, workflow systems will need to evolve and further embrace the new possibilities of these infrastructures. Also, not all research can be easily pipelined, for example exploratory research might be better carried out in an ad-hoc manner than with workflows and the overhead this implies. A Jupyter Notebook as used in in Demonstrator 1 or embedded in Galaxy (46) constitutes a promising way to make use of microservices for interactive analysis.

In summary, we showed that microservices allow for efficient horizontal scaling of analyses on multiple computational nodes, enabling the processing of large data sets. By applying a number of data (mzML (47), nmrML) and metadata standards (ISA serialisations for study descriptions (44, 45)), we also demonstrated a level of interoperability which has never been achieved in the context of metabolomics, by providing completely automated start-to-end analysis workflows for mass spectrometry and NMR data. The ability to instantiate VREs close to large datasets, such as on local servers within a hospital for Demonstrator 2, makes it possible to use the VRE on sensitive data that is not allowed to leave the current environment for ELSI reasons. While the current PhenoMeNal VRE implementation uses Docker for software containers and Kubernetes for container orchestration, the microservice methodology is general and not restricted to these frameworks. Likewise, the choice of Luigi and Galaxy was here used to demonstrate the capabilities of workflow management microservices in cloud environments. In fact, our microservice architecture accounts for other major workflow engines such as Nextflow (32) or Snakemake (48). Hence it is possible to use any of such workflow engines in our VRE and still produce reproducible results. In addition, despite some of our workflows were novel in the context of metabolomics (e.g. Demonstrator 2) and can be readily applied on other datasets, their main contribution in this work is to showcase scalability and interoperability of the microservices methodology. Finally, we emphasise that the presented methodology goes beyond metabolomics and can be applied to virtually any field, lowering the barriers for taking advantage of cloud infrastructures and opening up for large-scale integrative science.

## Acknowledgements

This research was supported by The European Commission’s Horizon 2020 programme funded under grant agreement number 654241 (PhenoMeNal), The Swedish Research Council FORMAS, Uppsala Berzelii Technology Centre for Neurodiagnostics and Åke Wiberg Foundation. We kindly acknowledge contributions by Daniel Jacob (INRA) and to cloud resources by SNIC Science Cloud, Embassy Cloud, and CityCloud. The funders had no role in study design, data collection and analysis, decision to publish, or preparation of the manuscript.

## Author contributions

KK, MAC, MC, PEK, SH contributed to Demonstrator 1. CR, KK, KP, PEK, SH, SN contributed to Demonstrator 2. KK designed the study in Demonstrator 2. JB performed collection of samples and characterization of the multiple sclerosis cohort. SH performed the mass spectrometry experiment in Demonstrator 2. DS, KP, PEK, PM, RMS, contributed to Demonstrator 3. AGB, CF, DJ, MCA, MVV, PDA, PM, PRS, SAS, TH and VS contributed to Demonstrator 4. GZ, LP, PEK and PM contributed to developments of Galaxy in Kubernetes. AL and MC contributed to the development of Luigi in Kubernetes. AL, MAC, MC and NS developed KubeNow. PM contributed to Galaxy-Kubernetes. EAT and PR contributed to containerizing of Workflow4Metabolomics tools. AGB, DJ, PRS and SAS contributed to ISA-API. DJ, EAT, KP, MVV, NS, OS, PEK, PM, PR, PRS, DS, RMS, RR and SB were involved in testing the containers and the VRE. PM, SIH and KH were involved in development and maintenance of the portal. MVV, PM and RMS contributed to the release. NK coordinated the PhenoMeNal project. CS conceived and managed the PhenoMeNal project. OS conceived and coordinated the study and e-infrastructure. All authors contributed to manuscript writing.

